# Brain inputs to the vestibular nuclei in lampreys

**DOI:** 10.64898/2026.05.06.723247

**Authors:** Cecilia Jiménez-López, Paula Rivas-Ramírez, Carmen Núñez-González, Marta Barandela, Manuel A. Pombal, Juan Pérez-Fernández

## Abstract

To avoid image blurring, the vestibulo-ocular (VOR) and the optokinetic (OKR) reflexes stabilize gaze. In all vertebrates, the VOR is mediated via direct projections from the vestibular nuclei to the motor nuclei that control the extraocular muscles. Lampreys show three vestibular nuclei that are well characterized in terms of their projections and sensory inputs, but much less is known about their inputs from other brain regions and the connectivity between them. Using tracer injections and electrophysiological recordings, we show that the lamprey vestibular nuclei are largely interconnected, while their inputs from other brain regions are scarce. The main rostral areas projecting to the vestibular nuclei are the pretectum and the ventral tier of the thalamus, which send ipsilateral inputs to the three vestibular nuclei.

## 1 Introduction

Animals constantly redirect their gaze to objects of interest in the visual scene. However, using the visual system for such advanced behaviors would not be possible without gaze stabilizing mechanisms to avoid image blurring. The VOR uses vestibular information to generate eye movements that counteract head rotations, and the OKR generates gaze stabilization by clamping the eyes to movements of the visual scene. These two reflexes are present in all vertebrate groups, and the neuronal mechanisms are conserved from lampreys (belonging to the oldest group of vertebrates) to mammals (Rovainen, 1976; Land, 2015; Wibble et al., 2022; Barandela et al., 2023). The VOR is mediated by a three-neuron pathway that conveys vestibular information to the motoneurons innervating the eye muscles. The labyrinth sends vestibular information to the vestibular nuclei, which in turn project directly to the ocular motor nuclei: the oculomotor (nIII), the trochlear (nIV) and the abducens (nVI) nuclei (Straka and Baker, 2013; Fritzsch, 2024). In lampreys, the vestibular nuclei are divided into three pairs of bilateral nuclei that are named according to their rostro-caudal location in the rhombencephalic alar plate, the anterior (AON), intermediate (ION) and posterior (PON) octavomotor nuclei and are located in the ventral nucleus of the OLA (González and Anadón, 1994; Pombal et al., 1994b). Additionally, in these animals, some primary vestibular fibers of the ascending branch are likely to contact motoneuron dendrites that belong to the three ocular motor nuclei, since nIII and nIV dendrites overlap with these primary octaval fibers (Fritzsch and Sonntag, 1988; Rodicio et al., 1992; Pombal et al., 1994a), and nVI motoneurons have been found to reach the ventral nucleus of the octavolateral area (OLA; Fritzsch and Collin, 1990; Pombal et al., 1994a; Pombal and Megias, 2019).

The AON is located close and dorsal to the trigeminal nerve entrance (Pflieger & Dubuc, 2000), and its neurons are large (>40 µm), round and spindle-shaped or bipolar. The AON projects to the nIII, mostly contralaterally although a few projections get to the ipsilateral nIII. Some AON neurons also have projections inside the ventral nucleus of the OLA, and other projections reach the contralateral third giant Müller cell in the caudal mesencephalon. Moreover, commissural projections from the AON cross the midline to target the contralateral AON (Pombal et al., 1994; Pombal et al., 1996; Wibble et al., 2022). The ION is located close to the entrance of the vestibular nerve (nVIII), and its caudal limit coincides with the level of the entrance of the posterior lateral line nerve (Bussières et al., 1999; Pflieger & Dubuc, 2000). In *Petromyzon marinus*, the ION contains around one hundred unipolar, bipolar and multipolar neurons, most of them between 20 and 40 µm of size (Bussières et al., 1999). The ION also sends projections to the nIII, mostly ipsilaterally (Pombal et al., 1994; Pombal et al., 1996). Axons leaving the ION also run caudally and medially to the ipsilateral spinal cord (Bussières et al., 1999), coursing together with the medial longitudinal fasciculus (MLF) through the medium rhombencephalic reticular nucleus (MRRN) and the posterior rhombencephalic reticular nucleus (PRRN) (Fritzsch et al., 1990; Bussières et al., 1999), while others reach the contralateral ION (Bussières et al., 1999). Fibers from both the AON and the ION have been also suggested to make monosynaptic contacts onto neurons of the nIV, although this remains to be proven (Pombal et al., 1994; Pombal et al., 1996). The PON is located caudal and slightly dorsal to the entrance of the posterior lateral line nerve (Pflieger & Dubuc, 2000). In *P. marinus* the PON contains around 70 neurons of medium size (20-40 µm) and large size (>40 µm). Most of them are unipolar, although bipolar and multipolar neurons were also reported (Bussières et al., 1999). Projections from the PON ran ventrally, cross the midline and then bifurcate so that a descending tract courses towards the spinal cord whereas the other, a short ascending tract, projects to the motoneurons of the nVI (Pombal et al., 1994; Pombal et al., 1996). The AON and the ION have been shown to receive also projections from the pretectum (PT), providing the basis for the OKR (Wibble et al., 2022), but whether pretectal projections reach the PON is not yet known.

Connectivity studies of the vestibular nuclei in lampreys have been thus focused mostly on their projections. However, although connections from the AON and the ION to their respective contralateral counterparts have been reported, the full set of vestibular connections is not available yet. Additionally, the inputs of the vestibular nuclei have been analyzed mostly in terms of the primary octaval fibers and lateral line nerves targeting them (Rubinson 1974; Northcutt, 1979; González and Anadón, 1994; Fritzsch et al., 1984; Ronan and Northcutt, 1987; González and Anadón, 1992), whereas inputs from other rostral brain regions have been only scarcely analyzed to the AON and the ION (Wibble et al., 2022). In the present study, the inputs from other brain regions to the AON, ION, and PON, as well as the interconnectivity between them were investigated by combining tracer injections with electrophysiological recordings. Our results show that these three nuclei are highly interconnected in lampreys, whereas they receive inputs from few other brain regions.

## 2 Results

The results obtained were consistent for the different species and stages studied (see Methods). Since no obvious differences were detected between specimens, the circuitry here described applies to all, although all the images showed in the Figures correspond to adult specimens. It is also remarkable that neurobiotin (low molecular weight) and dextran-rhodamine (high molecular weight) tracer injections (see Methods) lead to the same retrograde labeling throughout all the described results.

### 2.1 Inputs to the AON

The AON is the most well studied vestibular nucleus in lampreys, regarding both its outputs to motor areas and inputs from forebrain regions (see above). However, apart from its interconnectivity with the contralateral AON, no information exists about whether it receives projections from the other vestibular nuclei (Pombal et al., 1994; Pombal et al., 1996; Wibble et al., 2022). To further confirm previously described inputs, and to analyze the inputs to the AON from the other vestibular nuclei, we first performed tracer injections in this region (N=33) and analyzed the presence of retrogradely labeled cells (Figure 1A, inset). As previously described, the most rostral population that projects to the AON is located in the ventral aspect of the thalamus (Th; not to be confused with the old ventral thalamus, which is currently known as the prethalamus in lampreys, Pombal et el., 2009; Wibble et al., 2022). Retrogradely labeled cells were scarce and ipsilateral to the injection site (Figure 1A, arrow). Occasionally, neurons projecting to the AON were also found in more dorsal thalamic regions (not shown).

**Figure 1.**
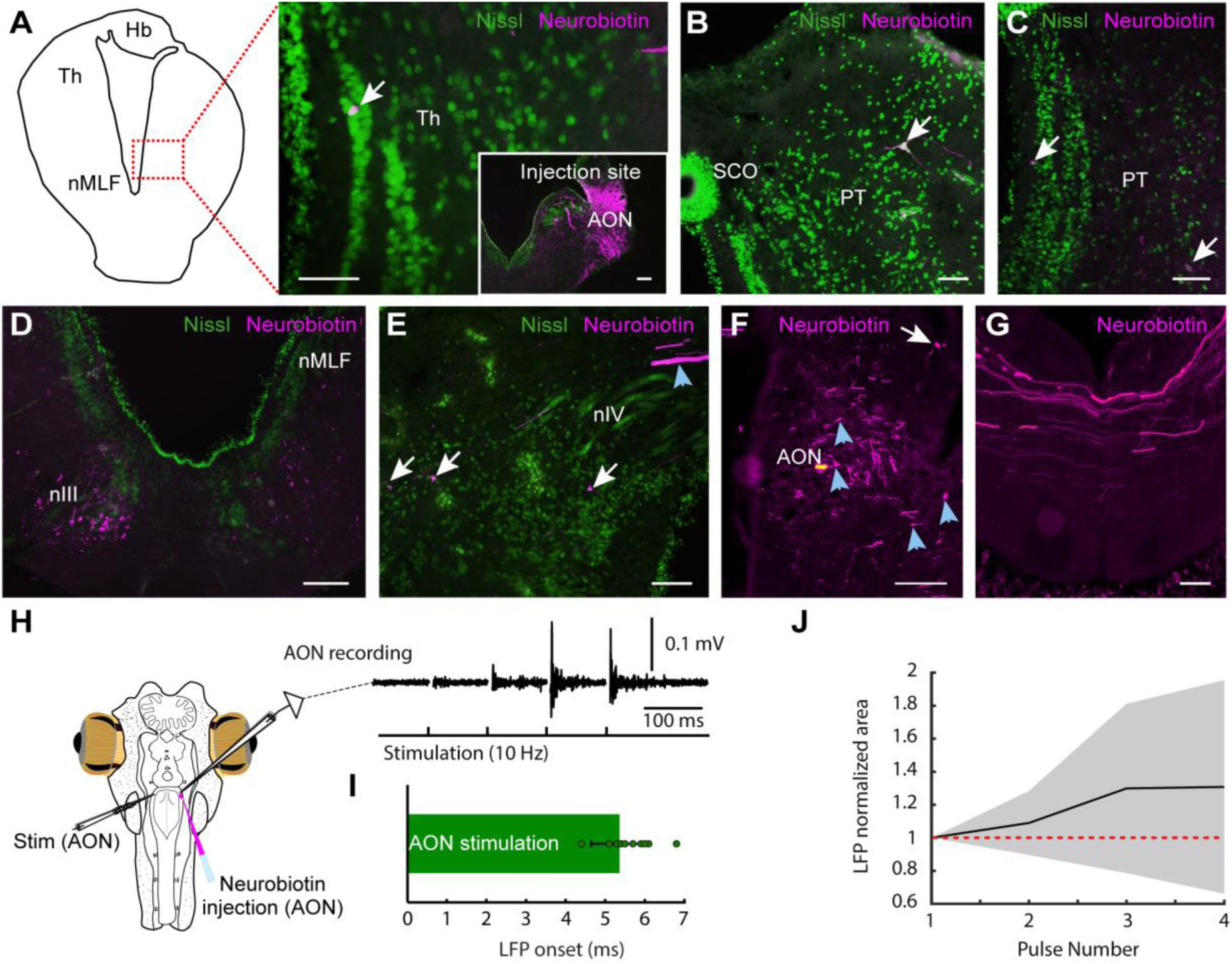
AON inputs from rostral brain areas and bilateral connections with the contralateral AON. (A) Micrograph (right) showing a retrogradely labeled neuron (arrow) in the ventral aspect of the ipsilateral thalamus (Th, the location is shown in the schematic at the left) after a neurobiotin injection in the anterior octavomotor nucleus (AON). The injection site is shown in the inset. (B) A retrogradely labeled neuron (arrow) in the lateral aspect of the ipsilateral pretectum (PT) at the level of the subcommissural organ (SCO), whose dendrites extend towards the region where the optic tract is located. In (C), two more AON projecting neurons are shown (arrows), in both medial and lateral aspects of the ipsilateral PT. (D) Anterogradely labeled fibers in the oculomotor nucleus (nIII) and the nucleus of the medial longitudinal fasciculus (nMLF) confirm the accuracy of the neurobiotin injection. Note that they are ticker and more numerous contralaterally. (E) AON projecting cells (arrows) in the contralateral isthmic region, near the location of the trochlear nucleus (nIV). Crossing primary vestibular afferents labeled due to passage in the AON region can also be seen at this level (blue arrowhead). (F) Retrogradely labeled neurons were found in the contralateral octavolateral area (arrow). Moreover, both numerous retrogradely labeled cells (blue arrowheads) and anterogradely labeled fibers are observed in the contralateral AON indicating that there is strong communication between the AON of both sides, and, accordingly, numerous fibers can be seen crossing at the ventral midline, as shown in (G). (H) Representative electrophysiological recording in the AON (right) in response to electric stimulation of the contralateral AON. The experimental preparation is shown in the schematic to the left. (I) Graph showing the onsets of the AON responses evoked by stimulation of the contralateral AON. (J) Mean responses in the AON in response to a 4 pulses electric stimulation of the contralateral AON (10 Hz). In the graphs, data are shown as mean ± s.d. Scale bars = 150 µm.

Retrogradely labeled cells were also confirmed in the PT, both close to the optic tract but also in more medial aspects, always ipsilateral to the injection site (Figures 1B, C, arrows; Wibble et al., 2022). In the mesencephalon, the accuracy of the tracer injections was confirmed by the presence of abundant anterogradely labeled thick fibers in the contralateral nIII, and less abundant fibers in the ipsilateral nIII (Figure 1D; Pombal et al., 1994; Pombal et al., 1996; Wibble et al., 2022; Barandela et al., 2023). No retrogradely labeled cells were found in mesencephalic areas, apart from occasional cells in the ipsilateral torus semicircularis (not shown). Interestingly, no retrogradely labeled cells were found in the optic tectum and the nucleus of the medial longitudinal fasciculus (nMLF), although numerous anterogradely labeled fibers were found in this last region (Figure 1D; Wibble et al., 2022). In the isthmic region of the rhombencephalon, abundant retrogradely labeled cells were found ipsilaterally, in the region of the nIV. Although it cannot be excluded that some of these cells were labeled because of their proximity to the injection site, retrogradely labeled neurons were also observed in the contralateral side (Figure 1E, arrows), indicating that neurons in this region project to the AON. At this level, thick primary vestibular afferents labeled due to passage in the AON region can also be seen (Figure 1E, arrowhead; Pombal and Megías, 2019).

Retrogradely labeled neurons were observed in the contralateral AON, as previously shown (Figure 1F, arrowheads; Pombal et al., 1996; Wibble et al., 2022), and numerous retrogradely and anterogradely labeled fibers crossing the ventral midline could be observed connecting both AON (Figure 1G). Retrogradely labeled neurons could be also observed in the contralateral OLA, close to the lateral line mechanoreceptor afferents of the medial nucleus (Figure 1F, arrow). To confirm that the AON influences its contralateral counterpart, we performed electrophysiological recordings in the AON (N=7; n=21) to analyze its activity in response to electrical stimulation of the contralateral AON (Figure 1H; 4 pulses, 10 Hz). As expected by the anatomical results, activity was evoked with a short latency (5.36±0.72 ms, Figures 1H, I), in agreement with a monosynaptic input (von Twickel et al., 2019). The evoked responses were potentiating (Figures 1H, J).

The AON also receives bilateral projections from the other two vestibular nuclei. Retrogradely labeled neurons were found both in the ipsilateral (Figure 2A, arrow) and the contralateral (Figure 2E, arrow) ION, after tracer injection into the right AON. Labeled cells were not numerous, but they were consistently found at different rostrocaudal levels in the ION. To confirm this connection, electrophysiological recordings were performed in the AON while the ipsilateral (N=4, n=12) or the contralateral (N=5, n=14) ION was electrically stimulated. In both cases, the evoked responses presented short latencies confirming that they are monosynaptic inputs (5.35±0.97 ms for ipsilateral ION; Figures 2B, C; and 5.54±0.86 ms for contralateral ION; Figures 2F, G) and were potentiating (Figure 2D for the ipsilateral ION and Figure 2H for the contralateral ION).

**Figure 2.**
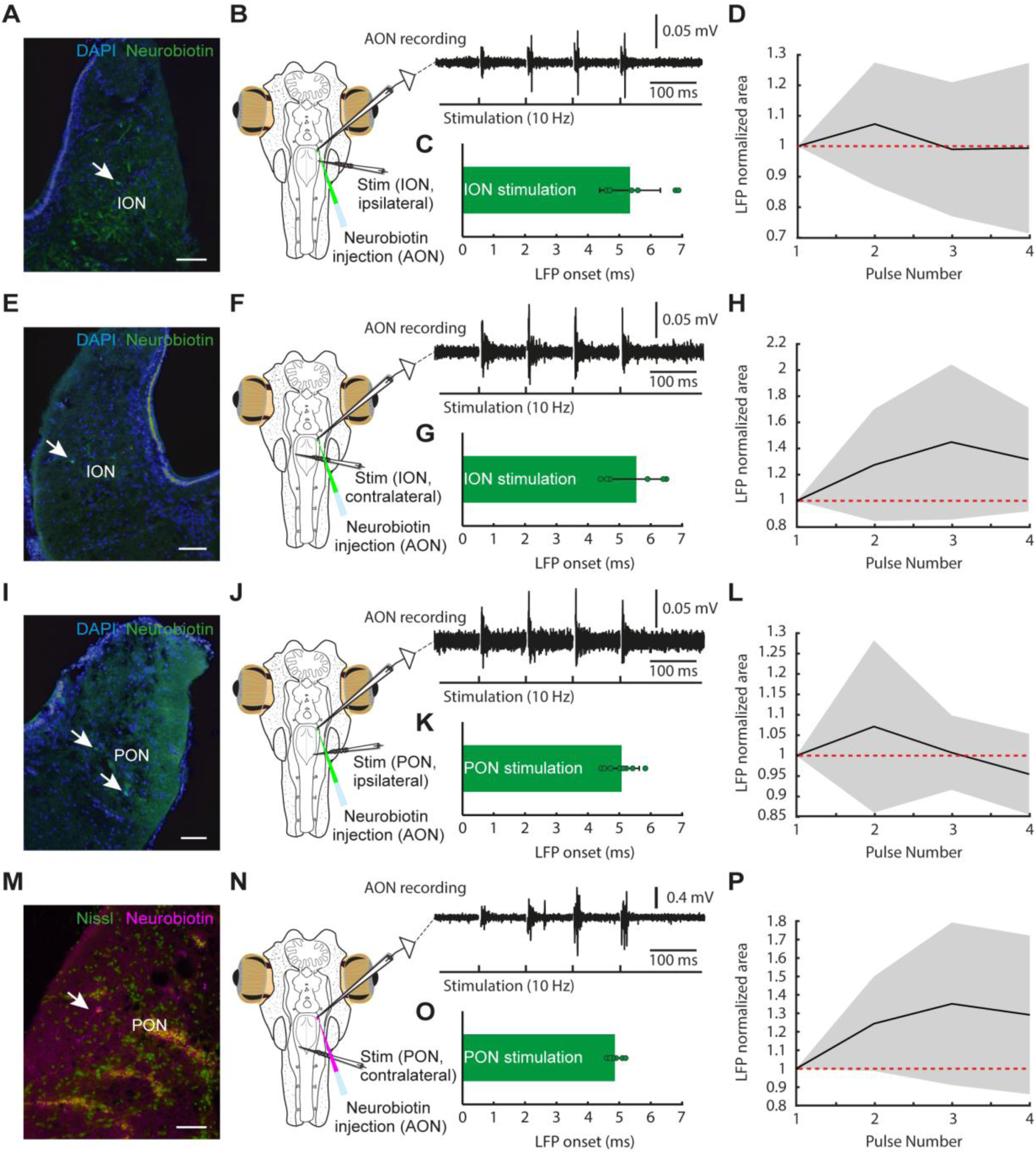
Bilateral AON inputs from the other two vestibular nuclei. (A) A representative retrogradely labeled neuron in the ipsilateral intermediate octavomotor nucleus (ION) after a neurobiotin injection in the anterior octavomotor nucleus (AON) is shown (arrow). (B) Electrophysiological recording in the AON in response to electric stimulation of the ipsilateral ION (4 pulses, 10 Hz). The experimental preparation is shown in the schematic to the left. (C) Graph showing the mean onset of the AON responses evoked by stimulation of the ipsilateral ION. The mean responses evoked is shown in the graph in (D). (E) Retrogradely labeled neurons could also be seen in the contralateral ION (arrow). In (F), an electrophysiological recording in the AON is shown in response to electric stimulation of the contralateral ION (4 pulses, 10 Hz). The experimental preparation is shown in the schematic to the left. The graph in (G) shows the mean onset of the evoked responses, and in (H) the mean responses for each of the 4 pulses is plotted. (I) Retrogradely labeled neurons in the posterior octavomotor nucleus (PON, arrows) after neurobiotin injection in the ipsilateral AON. (J) Evoked activity in the AON after a 4 pulses electrical stimulation of the ipsilateral PON (10 Hz). The experimental preparation is shown in the schematic to the left. (K) Plot showing the mean onset of the evoked responses after stimulation of the ipsilateral PON. (L) Graph showing the mean responses after the 4 pulses electrical stimulation of the ipsilateral PON. (M) Retrogradely labeled neurons were also observed in the contralateral PON (arrow). (N) Representative trace of the responses evoked in the AON after electrical stimulation of the contralateral PON (4 pulses, 10 Hz). The experimental preparation is shown in the schematic to the left. The mean onset of the evoked responses is shown in the graph in (O), and the mean responses after each of the 4 pulses in the graph in (P). In all the graphs, data are shown as mean ± s.d. Scale bars = 150 µm.

Finally, we also observed retrogradely labeled cells that were present in both the ipsi-(Figure 2I, arrows) and the contralateral (Figure 2M, arrow) PON. Electrical stimulation in both the ipsi- (N=4, n=12) and the contralateral (N=4, n=12) PON confirmed the monosynaptic excitatory connections from both nuclei to the AON (5.05±0.56 ms for the ipsilateral PON, Figures 2J, K; and 4.86±0.2 ms for the contralateral PON, Figures 2N, O) and showed that, as for the inputs from the other vestibular nuclei, the responses were potentiating (Figure 2L for the ipsilateral PON and Figure 2P for the contralateral PON). The results obtained from tracer injections and electrophysiology experiments show that the AON receives inputs from all the other ipsi- and contralateral vestibular nuclei, including the contralateral AON, while inputs from other brain regions are scarce.

### 2.2 Inputs to the ION

To characterize the projections from other brain areas to the ION, tracer injections were performed in this region (N=22; Figure 3A, inset). The most rostral population of retrogradely labeled cells was found in the thalamus, in the same location where the cells projecting to the AON were found. As for the AON, the number of thalamic cells that project to the ION is small and located ipsilaterally (Figure 3A, arrows). Retrogradely labeled cells were also found in the ipsilateral pretectum, as previously described (Figure 3B, arrow; Wibble et al., 2022). Unlike the AON, the cells projecting from PT to the ION were always located in medial aspects of the PT. As expected, anterogradely and bilaterally labeled fibers were observed in the nIII that were more numerous in the ipsilateral side, confirming the accuracy of the tracer injections (Figure 3C, arrows; Pombal et al., 1996). In the bilateral dorsal isthmic region, retrogradely labeled neurons were also observed, although they were less numerous than those observed after tracer injections in the AON, what could mean that the AON injections also affected some isthmic axons coursing towards the ION or the PON (Figure 3D, arrows). The isthmic neurons projecting to the ION were also located near the nIV.

**Figure 3.**
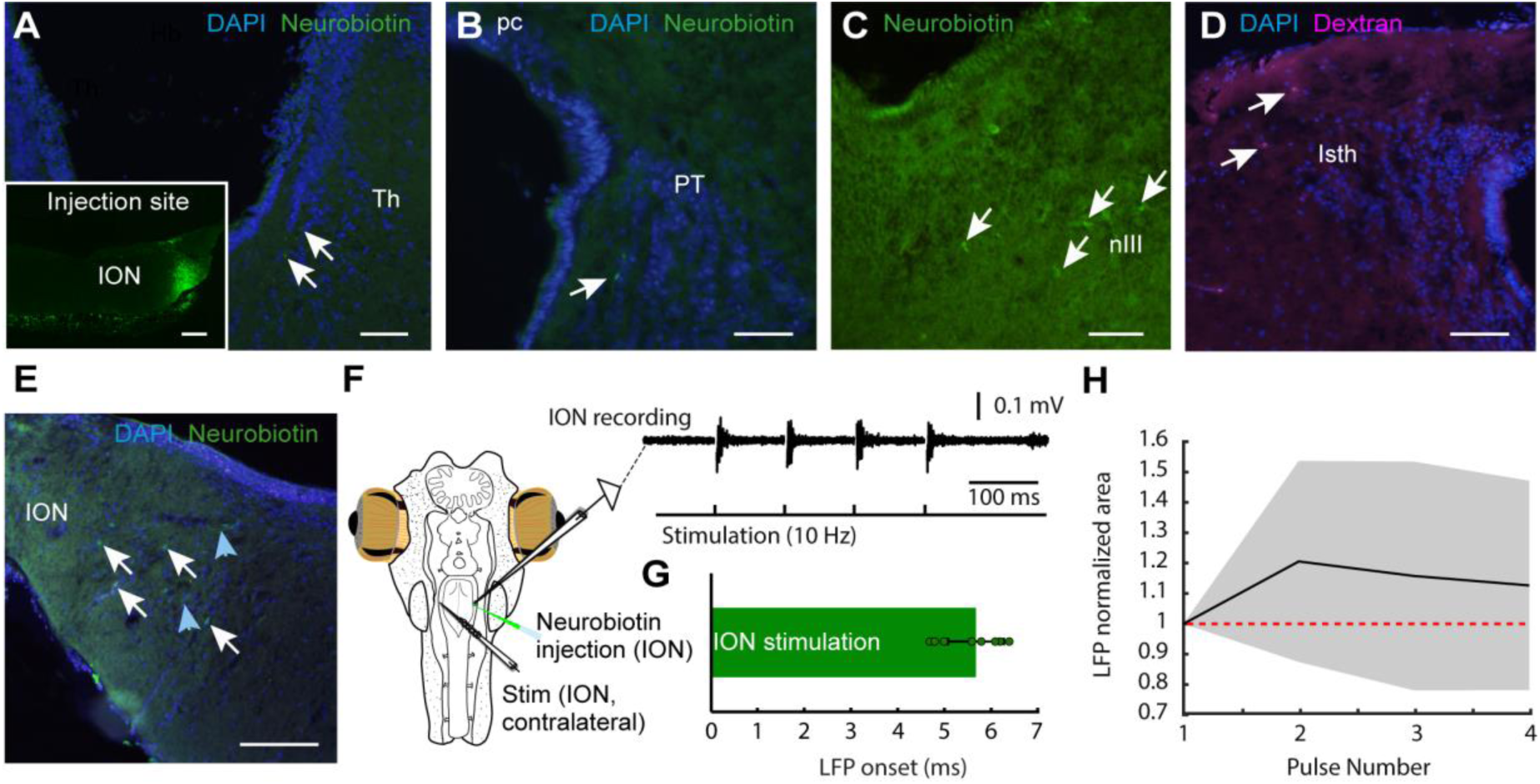
ION inputs from rostral brain areas and bilateral connections with the contralateral ION. (A) Retrogradely labeled neurons in the ventral aspect of the thalamus (Th; arrows) after neurobiotin injection in the ipsilateral intermediate octavomotor nucleus (ION). An image of the neurobiotin injection site is shown as an inset. (B) Retrogradely labeled neuron in the medial aspect of the ipsilateral pretectum (PT; arrow). (C) Anterogradely labeled fibers in the ipsilateral oculomotor nucleus (nIII; arrows) after a neurobiotin injection in the ION. (D) Retrogradely labeled neurons in the contralateral region of the isthmus (Isth; arrows). (E) Abundant retrogradely labeled cells (arrows) and anterogradely labeled fibers (blue arrowheads) were observed in the contralateral ION. (F) Stimulation of the contralateral ION (4 pulses, 10 Hz) resulted in activity in the ipsilateral ION. The experimental preparation is shown in the schematic to the left. (G) Graph showing the mean onset of the evoked activity in the ION after the stimulation of its contralateral counterpart. (H) Graph showing the mean responses evoked in the ION after each of the 4 pulses. In both graphs, data are shown as mean ± s.d. Abbreviations: pc, posterior commissure. Scale bars = 150 µm in A, B and D, 250 µm in A (inset) and E, 50 µm in C.

As in the case of the AON, tracer injections in the ION lead to retrogradely labeled neurons (arrows) and anterogradely labeled fibers (arrowheads) in its contralateral counterpart (Figure 3E). These results show that the ION also receives projections from the contralateral ION, as previously reported (Bussières et al., 1999; Wibble et al., 2022). Commissural fibers connecting both nuclei could be also observed crossing the ventral midline (both anterograde and retrogradely labeled), as well as retrogradely labeled neurons in the contralateral OLA, which are likely located in the limit of the medial nucleus (not shown). Electrical stimulation applied to the contralateral ION (N=5, n=15) resulted in short latency responses in the ION (5.69±0.6 ms, Figures 3F, G), corroborating the direct connection between both ION. The observed responses were potentiating (Figure 3H).

The ION also receives projections from the other octavomotor nuclei. It is important to note here that for ipsilateral projections, the results of these injections should be taken with caution. Since reciprocal connections exist between the AON and the PON (see above for AON inputs and below for PON inputs), it is possible that some of the retrogradely labeled cells in the AON and PON after injections in the ION could be due to fibers of passage. With this in mind, retrogradely labeled neurons were observed in the ipsi- (Figure 4A, arrows) and contralateral AON (Figure 4E, arrows), indicating the presence of connections from both anterior vestibular nuclei to the ION. To corroborate the projection from the ipsilateral AON, we electrically stimulated this nucleus (N=6, n=17) while recording in the ION (Figure 4B; 4 pulses, 10 Hz), which resulted in short latency responses confirming the monosynaptic inputs from the ipsilateral AON (5.78±0.8 ms, Figures 4B, C). The evoked responses were potentiating (Figure 4D). Electrophysiological recordings were also performed in the ION (N=7, n=21) while electrically stimulating the contralateral AON (Figure 4F; 4 pulses, 10 Hz). In this case, the monosynaptic inputs were also confirmed, according to their short latency (5.49±0.71 ms, Figures 4F, G), and the evoked responses were potentiating too (Figure 4H).

**Figure 4.**
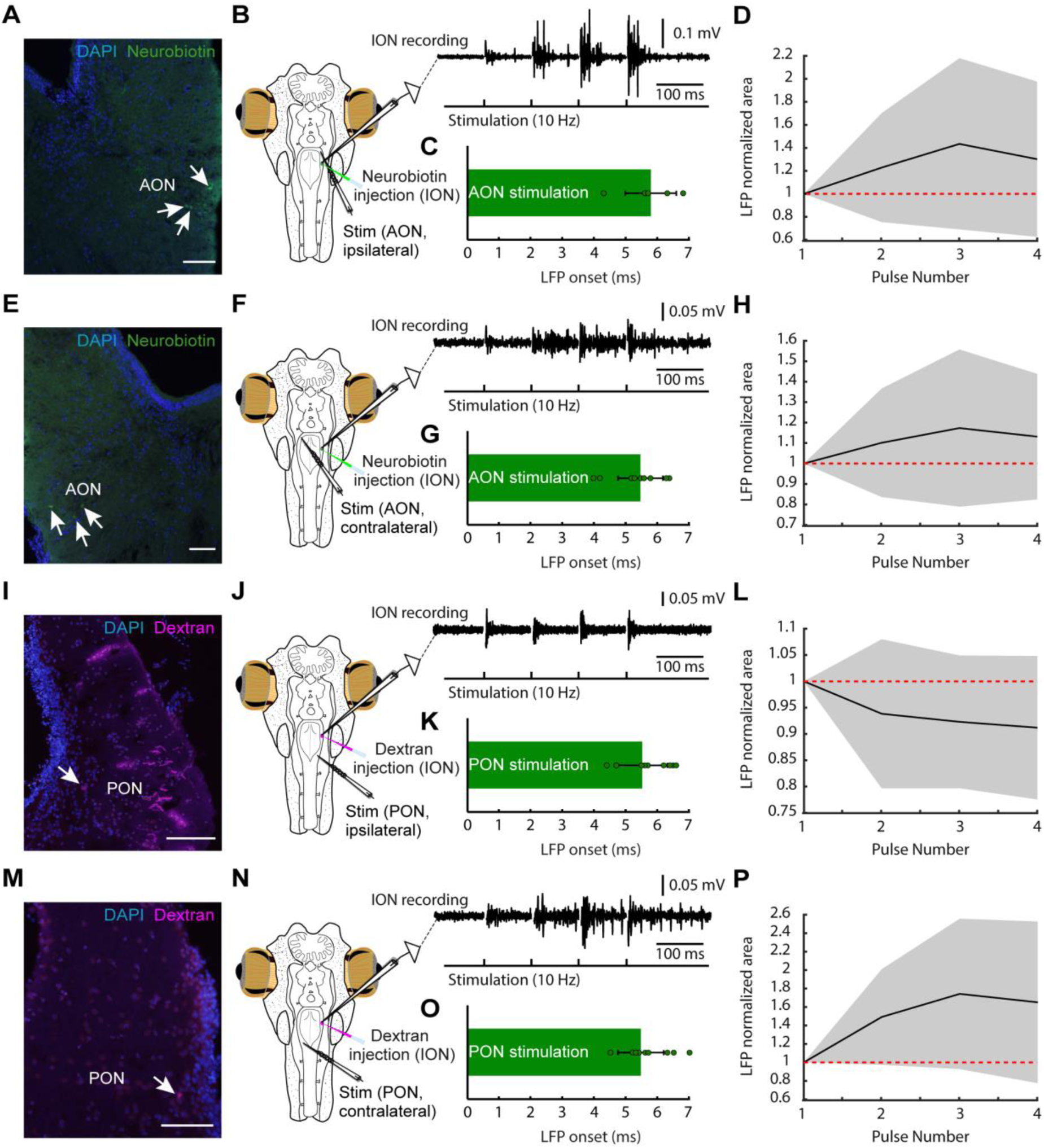
Bilateral ION inputs from the other two vestibular nuclei. (A) Photomicrograph showing retrogradely labeled neurons (arrows) in the ipsilateral anterior octavomotor nucleus (AON) after a neurobiotin injection in the intermediate octavomotor nucleus (ION). (B) Representative trace showing the activity evoked in the ION in response to electric stimulation of the ipsilateral AON (4 pulses, 10 Hz). The experimental preparation is shown in the schematic to the left. (C) Graph showing the mean onset of the ION responses evoked by stimulation of the ipsilateral AON. (D) Graph showing the mean responses evoked after each of the 4 stimulation pulses. (E) Retrogradely labeled neurons in the contralateral AON (arrows). (F) Electrophysiological recording in the ION in response to electric stimulation of the contralateral AON (4 pulses, 10 Hz). The experimental preparation is shown in the schematic to the left. The graph in (G) shows the mean onset of the evoked responses, and in (H) the mean responses for each of the 4 pulses is plotted. (I) Retrogradely labeled neuron in the posterior octavomotor nucleus (PON, arrow) after a dextran-rhodamine injection in the ipsilateral ION. (J) Trace showing the evoked activity in the ION after a 4 pulses electrical stimulation of the ipsilateral PON (10 Hz). The experimental preparation is shown in the schematic to the left. (K) Plot showing the onsets of the evoked responses after stimulation of the ipsilateral PON. (L) Graph showing the mean responses after each of the 4 pulses electrical stimulation of the ipsilateral PON. (M) Retrogradely labeled neurons were also observed in the contralateral PON (arrow). (N) Representative trace of the responses evoked in the ION after electrical stimulation of the contralateral PON (4 pulses, 10 Hz). The experimental preparation is shown in the schematic to the left. The mean onset of the evoked responses is shown in the graph in (O), and the mean responses after each of the 4 pulses in the graph in (P). In all the graphs, data are shown as mean ± s.d. Scale bars = 200 µm in A and M, 150 µm in E, 250 µm in I.

In both posterior octavomotor nuclei retrogradely labeled neurons were found, showing that also the PON in both sides projects to the ION (Figures 4I, M, arrows). To confirm that the ipsilateral PON sends information to the ION, electrical stimuli were applied to the ipsilateral PON (N=5, n=15) while recording the activity of ION neurons (4 pulses, 10 Hz). The activity was evoked with a short latency (5.52±0.8 ms, Figures 4J, K) demonstrating that inputs were monosynaptic. Interestingly, unlike all the other connections between vestibular nuclei, in this case the responses had an inhibitory nature (Figure 4L). Electrophysiological recording in the ION while the contralateral PON was stimulated (N=5, n=15) also resulted in responses with a short latency (5.46±0.72 ms, Figures 4N, O) verifying that the inputs are monosynaptic. Unlike the ipsilateral PON, the inputs from the contralateral PON were potentiating (Figure 4P).

### 2.3 Inputs to the PON

The PON is the less studied vestibular nucleus in lampreys in terms of the inputs that receives from other brain regions. Therefore, to characterize these projections to the PON, tracer injections were performed in this nucleus (N = 43; Figure 5A, inset). Our results confirmed that the ventral aspect of the thalamus also projects to the PON, constituting the most rostral population projecting to this vestibular nucleus. Retrogradely labeled cells were observed in the same location where the neurons projecting to the AON and the ION were found and, as in the case of the other vestibular nuclei, the number of cells that project to the PON is small and ipsilaterally located (Figure 5A, arrows). Retrogradely labeled cells were also found in the ipsilateral PT (Figure 5B). As for the AON, the cells projecting from the PT to the PON were located both in the lateral (arrows) and medial (arrowhead) aspect of the PT. In the dorsal isthmic region retrogradely labeled neurons were also observed near the bilateral nIV (Figure 5C, arrow).

**Figure 5.**
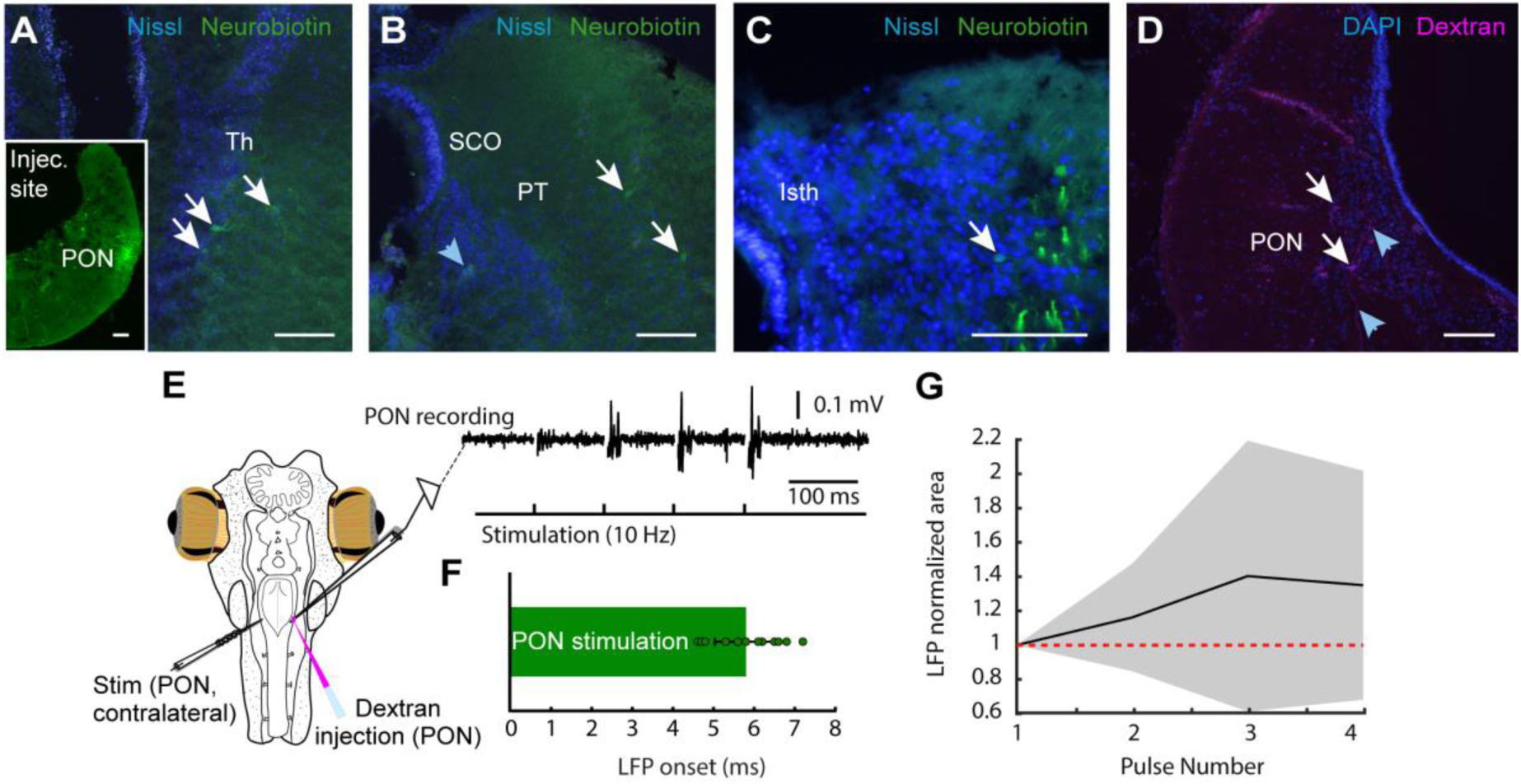
PON inputs from rostral brain areas and bilateral connections with the contralateral PON. (A) Retrogradely labeled neurons in the ventral aspect of the ipsilateral thalamus (Th; arrows) after a neurobiotin injection in the posterior octavomotor nucleus (PON). The injection site is shown in the inset. (B) Retrogradely labeled neurons were also found in the ipsilateral pretectum (PT), both in the medial aspect (blue arrowhead) and in more lateral regions (arrows). (C) Retrogradely labeled neuron in the region of the ipsilateral isthmus (Isth; arrow) after a neurobiotin injection. (D) Retrogradely labeled neurons in the PON after a dextran-rhodamine injection in the contralateral PON (arrows) and anterogradelly labeled fibers (blue arrowheads). (E) Electrophysiological recording in the PON showing the activity evoked in response to electric stimulation of the contralateral PON (4 pulses, 10 Hz). The experimental preparation is shown in the schematic to the left. (F) Graph showing the mean onset of the evoked responses. (G) Graph showing the mean responses after each of the 4 applied stimulation pulses. In both graphs, data are shown as mean ± s.d. Scale bars = 100 µm.

As for the AON and ION, the PON was reciprocally connected with the PON in the contralateral side. Traces injections in the PON resulted in retrogradely labeled cells (arrows) and anterogradely labeled fibers (arrowheads) in the contralateral PON (Figure 5D), and both types of labeled fibers were observed coursing between them after crossing at the ventral midline (not shown). Besides, retrogradely labeled neurons were found in the contralateral OLA, again close to the medial nucleus (not shown). Electrophysiological experiments were used to corroborate this connection between both PON. Electrical stimulation in the PON (N=6; n=17) resulted in responses in its contralateral counterpart with short latencies (5.8±0.77 ms, Figures 5E, F) that were potentiating (Figure 5G).

Regarding connections with the other vestibular nuclei, retrogradely labeled neurons appeared in the ipsilateral (Figure 6A, arrow) and contralateral (Figure 6E, arrow) AON when tracer injections were performed in the PON. To ensure that the PON receives inputs from the ipsilateral and contralateral AON, the latter were electrically stimulated while electrophysiological recordings were performed in the PON (Figure 6B for ipsilateral AON and Figure 6F for contralateral AON; 4 pulses, 10 Hz). The results showed that the PON receives monosynaptic inputs, showing a short latency, from the ipsilateral AON (N=6; n=18; 5.38±0.33 ms, Figure 6C) and from the contralateral AON (N=8; n=24; 5.32±0.58 ms, Figure 6G). In both cases, the responses evoked were slightly potentiating (Figure 6D for ipsilateral AON and Figure 6H for contralateral AON).

**Figure 6.**
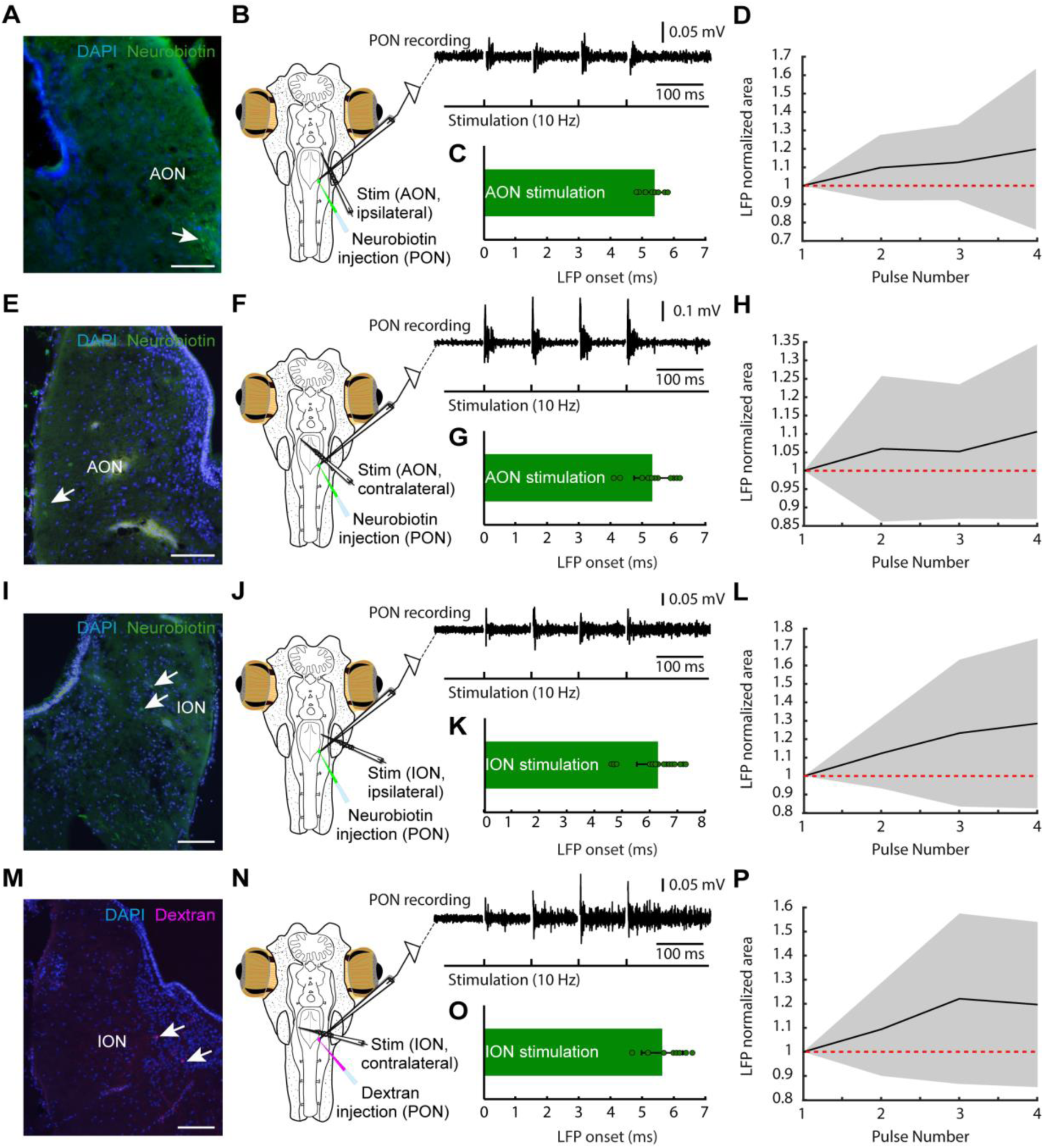
Bilateral PON inputs from the other two vestibular nuclei. (A) Photomicrograph showing a retrogradely labeled neuron (arrow) in the ipsilateral anterior octavomotor nucleus (AON) after a neurobiotin injection in the posterior octavomotor nucleus (PON). (B) Representative trace showing the activity evoked in the PON in response to electric stimulation of the ipsilateral AON (4 pulses, 10 Hz). The experimental preparation is shown in the schematic to the left. (C) Graph showing the mean onset of the PON responses evoked by stimulation of the ipsilateral AON. (D) Graph showing the mean responses evoked after each of the 4 stimulation pulses. (E) Retrogradely labeled neuron in the contralateral AON (arrow). (F) Electrophysiological recording in the PON in response to electric stimulation of the contralateral AON (4 pulses, 10 Hz). The experimental preparation is shown in the schematic to the left. The graph in (G) shows the mean onset of the evoked responses, and in (H) the mean responses for each of the 4 pulses is plotted. (I) Retrogradely labeled neurons in the intermediate octavomotor nucleus (ION, arrows) after a neurobiotin injection in the ipsilateral PON. (J) Trace showing the evoked activity in the ION after a 4 pulses electrical stimulation of the ipsilateral ION (10 Hz). The experimental preparation is shown in the schematic to the left. (K) Plot showing the mean onset of the evoked responses after stimulation of the ipsilateral ION. (L) Graph showing the mean responses after each of the 4 pulses electrical stimulation of the ipsilateral ION. (M) Retrogradely labeled neurons were also observed in the contralateral ION (arrows). (N) Representative trace of the responses evoked in the PON after electrical stimulation of the contralateral ION (4 pulses, 10 Hz). The experimental preparation is shown in the schematic to the left. The mean onset of the evoked responses is shown in the graph in (O), and the mean responses after each of the 4 pulses in the graph in (P). In all the graphs, data are shown as mean ± s.d. Scale bars = 100 µm.

Retrogradely labeled neurons were also found in the ipsi- (Figure 6I, arrows) and contralateral ION (Figure 6M, arrows). Electrophysiological recording in the PON confirmed the monosynaptic inputs from the ipsilateral ION showing short latencies (N=7; n=21; 6.29±0.76 ms, Figures 6J, K) and from the contralateral ION (N=5; n=15; 5.65±0.65 ms, Figures 6N, O). The electrical activation of the ipsilateral ION elicited a potentiating evoked response in the PON (Figure 6L), and the same is true when the contralateral ION was electrically activated (Figure 6P).

The results showed that the PON receives inputs from the other ipsi- and contralateral vestibular nuclei, including its contralateral counterpart, as well as from rostral brain areas that project to the most caudal vestibular nuclei of the lamprey, namely the ipsilateral ventral aspect of the thalamus, the ipsilateral PT and the bilateral dorsal isthmic region.

## 3 Discussion

Our anatomical and electrophysiological results show that the lamprey vestibular nuclei receive scarce inputs from other brain regions, whereas they are highly interconnected. Figure 7 summarizes the inputs from rostral brain regions and from the other vestibular nuclei to the AON (Figure 7A), to the ION (Figure 7B) and to the PON (Figure 7C).

**Figure 7.**
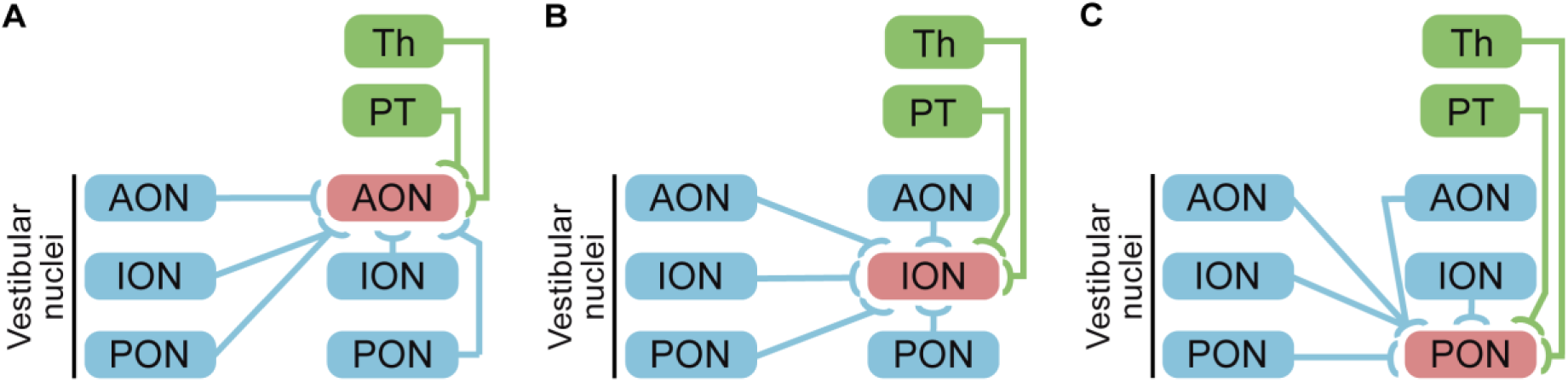
Summary of the main inputs from other brain regions to the lamprey vestibular nuclei. (A) Schematic showing the main rostral brain regions (green) projecting to the anterior octavomotor nucleus (AON, red), as well as the inputs the AON receives from the other vestibular nuclei (blue). (B) Schematic showing the main rostral brain regions (green) projecting to the intermediate octavomotor nucleus (ION, red), as well as the inputs the ION receives from the other vestibular nuclei (blue). (C) Schematic showing the main rostral brain regions (green) projecting to the posterior octavomotor nucleus (PON, red), as well as the inputs the PON receives from the other vestibular nuclei (blue). Abbreviations: Th, thalamus; PT, pretectum.

The most rostral brain region projecting to the lamprey vestibular nuclei was found to be the thalamus, specifically the ventral aspect of the Th, coinciding with previous data reported by Wibble et al., 2022, who demonstrated that the Th projects to the AON and ION. The results here exposed expand the circuitry, showing that the Th also projects to the PON, and indicating that the ventral aspect of the Th projects to the three ipsilateral vestibular nuclei. Interestingly, previous work reported a locomotor region, the diencephalic locomotor region (DLR), located in the ventral tier of the Th (El Manira et al., 1997; Ménard and Grillner, 2008). Given the importance of the vestibular nuclei for correct locomotion and the shared location of the DLR and the retrogradely labeled neurons that we found in the ventral aspect of the Th, one possibility is that the lamprey thalamic neurons projecting to the vestibular nuclei are part of this motor area and that their projections to the vestibular nuclei constitute efferent copies to convey information about the initiated movement.

Regarding the pretectum, their inputs to the AON and the ION had already been described (Wibble et al., 2022) but, again, we also show pretectal inputs to the most caudal vestibular nucleus, the PON. Interestingly, all the pretectal projections to the vestibular nuclei that we observed were ipsilateral, although the cell populations projecting to each vestibular nuclei turned out to be slightly different. Whereas injections in the AON and the PON resulted in retrogradely labeled neurons both in lateral (close to the optic tract), and medial aspects, ION projecting neurons were observed only in the medial aspect of the pretectum. The functional relevance of this difference remains to be tested, although the lack of retrogradely labeled neurons in the lateral aspect of the PT could be due to the restricted location of our ION injections. Therefore, we report that the three lamprey vestibular nuclei participate in visuo-vestibular integration, most likely both in gaze stabilization and body posture. Pretectal projections to the vestibular nuclei have been found in mammals, such as cats, rabbits and rats (Spence and Saint-Cyr, 1988; Watanabe et al., 2003; Hardcastle and Krapp, 2016), which reinforces the importance of this multisensory integration. Furthermore, the nucleus of the optic tract has been shown to enhance the gain of the angular VOR, indicating the active participation of the PT in modulating the VOR (Yakushin et al., 2000). This role in gaze stabilization is conserved in vertebrates, since inactivation of the PT also abolishes the OKR and its modulation of the VOR in lampreys (Wibble et al., 2022). Apart from this participation, the PT has been associated with optomotor reflexive responses that include spatial orientation contribution and postural regulation via projections to the reticulospinal neurons. Indeed, some explicit orientation and postural behaviors such as the dorsal light response (reaction to asymmetrical illumination) and negative phototaxis have been specifically related to the PT in lampreys. Besides, unilateral pretectal area lesions lead to perturbed postures, suggesting that the PT also influences posture control (Ullén et al., 1997). Therefore, although posture maintenance is highly based on vestibular information, it is strongly conditioned by visual integration in the PT, which likely mediates its motor commands both via direct projections to the reticular formation and indirectly by modulating the activity of the vestibular nuclei.

In the rostral limit of the rhombencephalon, in the dorsal isthmic region, retrogradely labeled cells were found close to the nIV. This region projects to the three vestibular nuclei both ipsi- and contralaterally. Previous work reported the location of higher-order brain areas that initiate locomotor activity in the lamprey, indicating that one of them is situated in the rostrolateral rhombencephalon (Paggett et al., 2004; Jackson and McClellan, 2011). Given the proximity of our labeled neurons to this region, one possibility is that they are part of this locomotor area, which likely sends locomotor-related information to the vestibular nuclei. On the other hand, although a functional cerebellum has not been demonstrated in lampreys, some authors have described a plate-like cerebellum in the rostrodorsal part of the lamprey rhombencephalon, in which transcription factors of the major cerebellar neurons (granule and Purkinje cells) have been detected (Sugahara et al., 2021). The retrogradely labeled neurons that we found were close to this region, suggesting that they could be part of this primordial cerebellum. Given the large communication between the vestibular nuclei and the cerebellum in other vertebrates (Barmack, 2023), and the high level of conservation of the vestibular system through evolution, it is reasonable that if the lamprey presented a cerebellum-like structure, it would be connected to the vestibular nuclei. Moving on to the possible nature of this labeled isthmic cells that we found, some of them are in the same region where ChAT immunoreactive neurons have been previously reported by Pombal et al. (2001), so it is also conceivable that these cells are cholinergic.

Since the lamprey neurons are not recovered by myelin, their synapses are slightly slower than those of other vertebrates, showing a latency of up to 8 ms for monosynaptic connections, versus the 1-5 ms showed by other vertebrates (von Twickel et al., 2019). The latencies that we observed were always between 4 and 6 ms, indicating that all connections between the lamprey vestibular nuclei reported are monosynaptic. Furthermore, neurobiotin (low molecular weight) and dextran-rhodamine (high molecular weight) injections lead to the same retrograde labeling, reinforcing that the involved synapsis are chemical rather than electrical, since neurobiotin can cross electrical synapses but dextran-rhodamine cannot (Song et al., 2016). The responses evoked in most cases were potentiating. The only exception was observed in the projections from the PON to the ipsilateral ION, whose responses showed an inhibitory nature. Both electrophysiological and anatomical experiments indicate that all the lamprey vestibular nuclei are interconnected.

In other vertebrates, vestibular nuclei are classically classified in four nuclei: the superior vestibular nucleus (SVN), located at the rostral part of the medulla; the lateral vestibular nucleus (LVN; also known as the Deiter’s nucleus), found in the caudal medulla; the medial vestibular nucleus (MVN), placed medially to the SVN; and the descending (also denominated inferior or spinal) vestibular nucleus (DVN), also located at the caudal part of the medulla (Brodal, 1974; Barmack, 2003). Additionally, the mammalian vestibular complex counts on some minor vestibular nuclei that also receive primary vestibular inputs, such as the parasolitary nucleus (Barmack et al., 1998), the Y-group (Frederickson and Trune, 1986; Blazquez et al., 2000) and the nucleus intercalatus (Brodal, 1984). Regarding their interconnections, in rabbits and cats it is known that the SVN is reciprocally connected with the DVN and MVN (Epema et al., 1988). Besides, the MVN sends projections to the DVN, and the SVN projects to both the ipsi- and contralateral Y-group. Finally, groups of larger neurons in the MVN, SVN and LVN receive inputs from smaller cells of MVN, SVN and DVN, but these connections are not reciprocal (Ito et al., 1985). Besides, connections of each vestibular nucleus with their contralateral counterpart have been described in the gerbil, except for the LVN and the parasolitary nucleus. Other contralateral projections found in this animal include the MVN projecting to the other 3 vestibular nuclei, and the SVN projecting to the LVN and the DVN (Newlands et al., 1989). In frogs, commissural projections between contralateral second order vestibular neurons have been demonstrated (Holler and Straka, 2001; Lambert and Straka, 2012). In fact, a commissural exchange of information between contralateral vestibular nuclei has been reported in many vertebrates, such as lizard (Richter et al., 1975), guinea pig (Babalian et al., 1997) and monkey (McCrea et al., 1987), but the details of these commissural projections have not been fully defined. Moreover, a recent study in mice showed that vestibular nuclei glutamatergic neurons receive inputs from other second order vestibular cells (Shi et al., 2021). Therefore, as in lampreys, the vestibular nuclei of other vertebrates are also interconnected, indicating that these connections are conserved in vertebrate evolution.

In lampreys the only known vestibular internuclear connections before the present study were the contralateral projections from AON to AON and from ION to ION (Pombal et al., 1994b, 1996; Bussières et al., 1999; Wibble et al., 2022). Now, we report a full map of the vestibular nuclei interconnections, showing that all the vestibular nuclei are connected among them. However, it is worth mentioning that, as in the rostral brain regions, the number of retrogradely labeled cells found to project to the other vestibular nuclei is low. Interestingly, the only exception to this are the projections between contralateral counterparts, suggesting a major communication between symmetrical vestibular information. In other vertebrates it is known that the vestibular nuclei have a push-pull system, so that when the head turns to the right, the vestibular system in that side is excited. This increased firing on the right not only generates motor commands for gaze stabilization and gaze control but also sends inhibitory signals to the left vestibular nuclei, further decreasing their already diminished activity. This inhibition has been shown to be both direct and feedforward, and excitatory commissural projections have been also demonstrated (Shimazu and Precht, 1966; Kong et al., 2024). It is clear that excitatory projections communicate all the vestibular nuclei in lampreys, since we could extracellularly record evoked activity. This suggests that, if inhibition is also present, this is likely feedforward. Interestingly, the retrogradely labeled neurons that we observed generally showed a small size. Tracer injections in the nIII have been reported to result in both large (referred to as octavomotor neurons) and small (referred to as ventral nucleus cells) retrogradely labeled neurons in the AON (Pombal et al., 1996). It is possible that the neurons that project to other vestibular nuclei correspond to the latter, although it is also possible that the neurons that communicate with the other vestibular nuclei are not the same that send the motor commands underlying the VOR; therefore, double tracer injections are needed to clarify this question.

In conclusion, our results show that the lamprey vestibular nuclei are all interconnected, highlighting that commissural connections between contralateral counterparts are the most abundant ones. Additionally, the three vestibular nuclei receive similar but scarce inputs from other brain areas.

## 4 Materials and Methods

### 4.1 Animal Model

Experiments were performed on 49 lampreys: 23 adult river lampreys (*Lampetra fluviatilis*), 8 adult, 4 postmetamorphic and 14 larval sea lampreys (*Petromyzon marinus*). The larvae were captured in the Furnia river (41°59’33.0 “N 8°41’33.0 “W, Galicia, NW Spain), the adult and postmetamorphic sea lampreys were obtained from local authorized commercial distributors, and adult river lampreys were kindly donated by Professor Sten Grillner (Karolinska Institutet, Sweden). All animals were kept in aquaria with continuous oxygenation and filtration. The experimental procedures were approved by Xunta de Galicia and the University of Vigo Committee for Animal use in Laboratory in accordance with the European Parliament directive (2010/63/EU) and the Spanish regulation on the protection of animals use for scientific purposes (RD 53/2013). The experiments were designed trying to minimize suffering and reducing the number of lampreys used while maximizing the data obtained.

### 4.2 Experimental procedures

Animals were deeply anesthetized with tricaine methane sulfonate dissolved in water (MS-222;100 mg/L; Sigma-Aldrich) before decapitation performed between the third and fourth gill openings. The dissection was done in a chamber containing refrigerated artificial cerebrospinal fluid (aCSF) with the following composition (in mM): 125 NaCl, 2.5 KCl, 2 CaCl_2_, 1 MgCl_2_, 10 glucose, and 25 NaHCO_3_, saturated with 95 % O_2_/5 % CO_2_ (vol/vol). To expose the brain and avoid muscle contractions during electrophysiological experiments, all muscles, viscera and skin were removed. The preparation was placed in another chamber perfused with aCSF to execute electrophysiological experiments and tracer injections.

### 4.3 Electrophysiological experiments

Extracellular recordings were performed to study the connectivity between vestibular nuclei. Tungsten microelectrodes (∼1–5 MΩ), connected to a differential AC amplifier (model 1700, A-M systems), were used to record extracellular activity in one vestibular nucleus while applying electrical stimulation to the other vestibular nuclei using a silver wire inside borosilicate micropipettes (od = 1.5 mm, id = 1.17 mm; Hilgenberg) filled with aCSF. Micromanipulators (model M-3333, Narishige) were employed to place the microelectrodes and micropipettes in the desired vestibular nuclei in each experiment. Ocular movements evoked by electrical stimulation of the different vestibular nuclei were used as a control to ensure that the stimulation was performed in the right location. This strategy was also used as a control for the injections of neuronal traces (see below). Recorded signals were digitized at 20 kHz using the pClamp 10.4 software. The same software was used to apply stimulation trains consisting of 4 pulses (10 ms duration pulses, 10 Hz) via a stimulus isolation unit (MI401, Zoological Institute, University of Cologne) used to control the stimulation intensity set (from 0.01 to 1 mA) to evoke neuronal activity.

### 4.4 Anatomical tract tracing

Two neuronal tracers were used to analyze the connectivity of the vestibular nuclei, neurobiotin (322,8 DA Vector Laboratories) and rhodamine-dextran (tetramethylrhodamine, 10,000 Da, Invitrogen). These tracers were separately dissolved in aCSF and injected into the different vestibular nuclei using a glass micropipette (borosilicate; od = 1. 5 mm, id = 1.17 mm; Hilgenberg) with a tip diameter of 10-20 μm connected to a pressurized air supply, attached to a micromanipulator (model M-3333, Narishige) to deliver the tracers in a precise manner. The brains were then kept in aCSF at 4 °C for 1 to 4 days in darkness to allow the transport. Then, the brains were fixed in 4 % formaldehyde and 14 % saturated picric acid in 0.1 M phosphate buffer (PB), pH 7.4, for 12-24 h, and cryoprotected in 20-40 % (wt/vol) sucrose in 1x phosphate-buffered saline (PBS) for 4-12 h. After this time, brains were embedded in OCT compound (Tissue-Tek, Sakura) and 30 μm thick transverse sections were obtained using a cryostat (Leica cm1950) and collected on gelatinized slides. Dextran-rhodamine-applied samples were directly mounted in DAPI-containing glycerol (DAPI Fluoromount-G, SouthernBiotech), as this tracer already comprises a fluorophore. To detect neurobiotin, the brain sections were incubated with Cy2-conjugated streptavidin (Alexa Fluor 488, 10,000 Da, Invitrogen, D22910) or Cy3-conjugated streptavidin (Jackson ImmunoResearch) 1:1000 in blocking solution (1 % bovine serum albumin, 10 % sheep serum, 0.1 % sodium azide and 0.3 % Triton X-100 in PB) in a dark humid chamber for 2 h, and then washed three times with PBS for 10 min each. After, the slides revealed with Cy2 were mounted in DAPI-containing glycerol (DAPI Fluoromount-G, SouthernBiotech), and those revealed using Cy3 were also incubated with Nissl stain (NeuroTrace 500/525 green fluorescent Nissl stain, Invitrogen) 1:500 and mounted in glycerol (Panreac Química S.L.U.).

### 4.5 Image analysis

All photomicrographs were obtained either with a digital camera (Nikon DS-Ri2) coupled to a Nikon ECLIPSE Ni-E microscope and to a Nikon Intensilight C-HGFI fluorescence source, where the acquisition software was NIS Elements D; or with a digital camera (Olympus DP71) coupled to an Olympus DX51 microscope and to an Olympus U-RFL-T fluorescence source, where the acquisition software was Olympus DP Controller. Then, exclusively the brightness and contrast of the photomicrographs were adjusted using ImageJ 1.53k and GIMP 2.1. Figure compositions were made using Adobe Illustrator CC 2019.

### 4.6 Quantification and statistical analysis

To determine whether neuronal activity was potentiating or inhibitory in response to 4 pulses electrical stimulation, the area under the curve after each applied pulse was measured using custom written functions in Matlab R2023b. Then, the area data were normalized to the first pulse. To explore the monosynaptic nature of the recorded extracellular responses, onset latencies were obtained by calculating the time from stimulus application to the first spike of the neuronal response. The number of animals used (N) and the number of experiments performed (n) were indicated throughout this document when applicable. Throughout the text and Figures, sample statistics were expressed as means ± s.d.

## 6 Author Contributions

Conceptualization: CJL, PRR, JPF; Experimental design: CJL, PRR, MP, JPF; Data acquisition: CJL, PRR, CNG, MB, MP; Data analysis: CJL, PRR, CNG, MB, MP, JPF; Writing: CJL, PRR, MP and JPF with inputs from all the authors; Supervision and Funding acquisition: JPF.

## 8 Acknowledgments

We are grateful to Manuel Megías for its constant technical and content support, and to Sten Grillner, Esra Öncel and the river station of A Freixa for helping with lamprey supply.

## 9 Data Availability Statement

The raw datasets for this study are available on request by contacting the corresponding author.

